# Power Analysis Provides Bounds for Genetic Architecture and Insights to Challenges for Rare Variant Association Studies

**DOI:** 10.1101/100891

**Authors:** Andriy Derkach, Haoyu Zhang, Nilanjan Chatterjee

**Affiliations:** Division of Cancer Epidemiology and Genetics, National Cancer Institute, National Institutes of Health, Rockville MD 20850; Department of Biostatistics, Bloomberg School of Public Health, and Department of Oncology, School of Medicine, John Hopkins University, Baltimore, MD 21205

**Author notes:** Corresponding author Nilanjan Chatterjee, Ph.D. Department of Biostatistics, Bloomberg School of Public Health, Johns Hopkins University, 615 N Wolfe St, Baltimore, MD 21205.

## Abstract

Genome-wide association studies are now shifting focus from analysis of common to uncommon and rare variants with an anticipation to explain additional heritability of complex traits. As power for association testing for individual rare variants may often be low, various aggregate level association tests have been proposed to detect genetic loci that may contain clusters of susceptibility variants. Typically, power calculations for such tests require specification of large number of parameters, including effect sizes and allele frequencies of individual variants, making them difficult to use in practice. In this report, we approximate power to varying degree of accuracy using a smaller number of key parameters, including the total genetic variance explained by multiple variants within a locus. We perform extensive simulation studies to assess the accuracy of the proposed approximations in realistic settings. Using the simplified power calculation methods, we then develop an analytic framework to obtain bounds on genetic architecture of an underlying trait given results from a genome-wide study and observe important implications for the completely lack of or limited number of findings in many currently reported studies. Finally, we provide insights into the required quality of annotation/functional information for identification of likely causal variants to make meaningful improvement in power of subsequent association tests. A shiny application, *Power Analysis for GEnetic AssociatioN Tests (PAGEANT)*, in R implementing the methods is made publicly available.

## Introduction

Over the last decade, genome-wide association studies (GWAS) of common variants of increasingly large sample sizes have been the main driving force for discovery of susceptibility loci associated with complex diseases and traits. While analysis of heritability suggests that common variants have further ability to explain additional variation of these traits ^1-11^, the focus of the field is inevitably shifting towards studies of less common and rare variants with the rapidly decreasing cost of sequencing technologies and increasing sophistication of imputation algorithms ^12-14^. However, limited or lack of findings from early studies ^15-29^ indicate that effect sizes of rare susceptibility variants in general are likely to be modest and discovery of underlying loci will require large sample size in future studies^30-32^.

Testing of associations at the level of genetic loci or regions using various aggregate-level statistics have been proposed as a strategy to improve power of discovery in association studies of rare variants^33-44^. Simulation studies have been used under various anticipated genetic architecture of the traits for the demonstration of potential power of these procedures ^35; 36; 44^. In particular, analysis of power for variance component based tests, such as the popular SKAT method, can be complex as they require specification of many different parameters including number of genetic variants under study, proportion of causal variants, allele frequency and effect-size distributions. Use of various functional and annotation information to identify likely pathogenic variants a priori has also been proposed as a strategy to improve power of rare variant association tests^45; 46^. To the best of our knowledge, however, there has been no systematic study of the effect of the use such extraneous information on power of the association tests.

In this report, we first describe approximations that allow analytic characterizations of power for popular aggregate-level association tests based on a few key parameters – thus dramatically reducing the complexity of power calculations. We perform simulation studies using allele frequency distribution observed in Exome Aggregation Consortium (ExAC)^47^ under various models for effect-size distributions to assess the accuracy of the proposed approximations in realistic settings. We then develop a framework for genome-wide power calculations based on underlying genetic architecture of a trait characterized by number of underlying causal loci and total variability they explain. We assess the power of a number of recently reported association studies of rare variants using the proposed framework and provide insights into the implications for lack of discoveries on bounds of genetic architecture of the underlying traits.

We also use the proposed framework to characterize power of association tests that may pre-select variants based on prior functional/annotation information. These derivations provide important insights into the required quality of annotation/functional information for identification of likely causal variants to make meaningful improvement in power of subsequent association tests. Finally, to facilitate convenient and rapid power calculations for rare variant association tests, we make a shiny app PAGENAT (Power Analysis for GEnetic AssociatioN Test) available in R.

## Material and Methods

### Existing Power Calculations

A variety of statistics have been proposed for testing genetic associations at the levels of genetic loci or regions by aggregating association statistics over multiple genetic variants ^33-44^. Multiple studies ^30; 42; 48^ have shown that existing methods can be classified as sum-based tests ^33; 34; 38; 43^, variance component tests ^35; 44; 48^, and hybrid tests that are functions of both classes ^36; 41; 42^. Here, we focus on sum-based and variance component tests. We do not consider hybrid tests because their power is usually close to one of the two components. Sum-based tests aggregate variant level association statistics by a linear combination in the forms

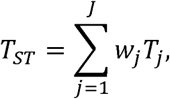

and variance component tests aggregate by quadratic combination in the forms

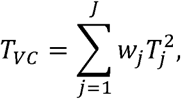

where *w*_*j*_s are weights that depend on MAF(*p*_*j*_) and *T*_*j*_s are score statistics for associations for individual SNPs (*j*=1,…,*J*), the latter of which are typically derived from a regression model, such as the linear or logistic regression ^36; 42; 48^.

Existing analytic power formulas for sum-based and variance component tests are complex functions of many parameters including number of genetic variants under study, proportion of causal variants, allele frequencies and effect-size distributions ^48^. The analytic power of a single-variant statistic

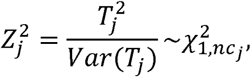

can be derived based on one degree-of-freedom chi-square distribution with a non-centrality parameter of the form 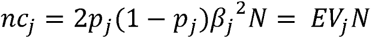, which depends on three parameters: effective sample size (*N*), level of the test (*α*) and proportion of phenotypic variation explained by the *j*^*th*^ variant (*EV*_*j*_)^4^.

Derkach et al.^48^ showed that analytic power for a sum-based test statistic *Z*_*ST*_,

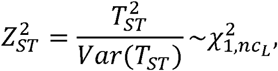

depends on the non-centrality 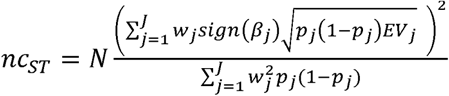 parameter which depends coefficients of explained variations associated with individual variants. Previous studies^35; 48^ have shown that a variance component statistic is asymptotically distributed as a linear combination of non-central chi-square random variables,

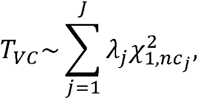

with non-centrality parameters *nc*_*j*_ = *EV*_*j*_*N* and weights *λ*_*j*_ = *W*_*j*_*P*_*j*_(1-*P*_*j*_)*N*. It has been suggested that analytic power calculations for variance component tests be done by approximating the asymptotic distribution of *T*_*VC*_ by a single non-central chi-square distribution matched up to four cumulants^49^. There test are also several modifications of this method matching higher moments to improve the tail probability approximation ^35; 50^; however, power differences seem to be marginal. The cumulants *c*_*k*_ of the statistic *T*_*VC*_ can be written as

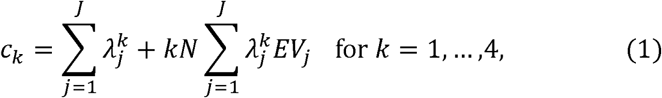

which require specification of effect-sizes (*EV*_*j*_) and allele frequencies of individual variants. The power calculations for aggregate level tests requiring specification of MAFs and genetic effects for individual variants have been implemented in several statistical packages ^35; 50; 51^.

### Approximate Power Calculations for Aggregate Tests

Following, we describe simple formulae for approximating power for different aggregate level tests using a limited number of key parameters. First, we show that for the sum-based test, under an assumption of independence between coefficients of explained variations and MAFs, we can roughly estimate non-centrality parameter as

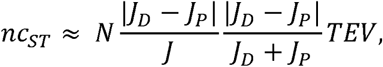

where *J*_*D*_ and *J*_*P*_ are numbers of deleterious and protective variants in a locus and 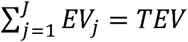 is the total proportion of variation explained by all variants in the locus (see Appendix A). If all of the causal variants in a locus are deleterious (or protective), then the non-centrality parameter can be characterized by *TEV* and the proportion of causal variants.

Next we consider simplified power calculations for variance component tests *T*_*VC*_ by approximating cumulants *C*_*k*_ in (1) as a function of the total proportion of variance explained by all variants within a locus (*TEV*). For example, if we assume independence between MAF and proportion of variation explained across individual variants, we can obtain a first-order approximation in the form

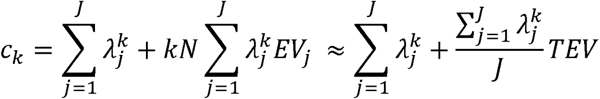

In Appendix B, we derive approximations of *C*_*k*_ for three commonly assumed relationships between genetic effects and MAFs and summarize them in Table S1. The first order approximations, which implicitly treat all variants in a locus to be causal, can be inaccurate when number of true causal variants is small. To improve accuracy, we propose second order approximations to estimate the sum 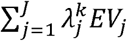 in (1) as function of *TEV* and number of underlying casual variants (*J*_*c*_) (see Appendix B and Table S1).

For example, if we assume the same hypothesis of independence between proportion of variation explained and MAF across individual variants, then we approximate 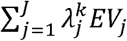 as 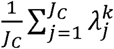

### Genome-wide power calculations and bounds on genetic architecture

Using the proposed power calculation framework, we further develop a mathematical framework to study bounds on genetic architecture of underlying traits from results reported in a genome-wide association study. We first characterize probability of number of discoveries in a given study as a function of sample size *N*, number of underlying causal loci *K* and the distribution of their effect-sizes (*TEV*). In Appendix C we show that probability of *M* discoveries in a study is

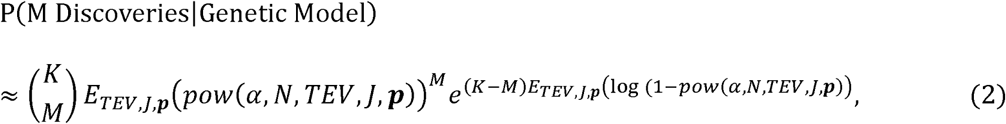

which depends on average power of the underlying association tests over the different causal loci in the genome and can be calculated by specifying number of underlying causal loci and distributions of proportion of phenotypic variations they explain (*TEV*), number of SNPs within a locus (*J*) and minor allele frequency ***p*** across the causal loci (see Appendix C).

Now, if M=m is the number of discoveries reported based in a given GWAS of sample size *N*, we can calculate P(M ≤ m) using the above formula based on empirical distributions for MAFs (***p***_*k*_), and size of genes (*J*_*k*_) observed in real data and various hypothesized values for number of causal loci (K) and parameters for underlying effect-size distributions for *TEV*. Specifically, we generated a class of L-shaped effect-size distribution using a two-parameter gamma distribution: Gamma(α, γ) with α ≤ 1 and restriction. Under this model, the total variance of a trait explained by causal loci is given *GEV* = *K*μ by μ ≈ αγ where. For various combination of K and μ, we evaluate the maximum value of P(M ≤ m) over different values of the dispersion parameter (α/γ = α^2^/μ) determined from wide range of possible values of α.

When this probability is low (e.g. < 5%), we conclude that the underlying model for genetic architecture is unlikely. For example, many recent studies have reported no discoveries based on gene-level association tests. In these studies, the probability of no discoveries, m=0 can be used to provide bound on genetic architecture of the underlying trait.

### Effects of filtering variants by extraneous information

We use the proposed framework to study the effects of filtering of variants based on prior functional/annotation information on the power of association tests. Here, power of association tests can be summarized as a function of sensitivity and specificity of the underlying filtering method. Sensitivity (Se) is the probability of selecting a variant given that it is truly causal, while specificity (Sp) is the probability of filtering a variant out given that is non-causal. If selection/filtering is independent of MAFs and proportions of variations explained, then the number of remaining variants after filtering in a locus is *J*_*s*_ *= Se. J*_*C*_ + (1-*SP*)·(*J*-*J*_*c*_) and the proportion of variation explained by them is *TEV*_*s*_ *= Se · TEV*. Now, with new values of and *J* and *TEV*, we estimate power for aggregated tests and compare them to corresponding base values if no filtering was applied (*e.g. Se* = 100%, but specificity is *Sp* = 0%).

If only a small subset of variants is selected, then sensitivity may be reduced as some true causal variants could be missed while specificity may improve because of removal of non-causal variants. If one takes random subset of the variants, then *Se* = 1-*Sp* as the casual and non-causal variants are selected at the same rate. If the functional/annotation information used for screening is predictive of whether the SNPs are likely to be causal for the trait of interest, then one would expect specificity > 1-sensitivity. Using the proposed framework, we explore power of aggregated tests for various combinations of sensitivity and specificity of the underlying filtering algorithm. Further, assuming than an underlying normally distributed continuous score represents the functional/annotation information for the SNPs, we evaluate receiver operating characteristic (ROC) curves generated by combination of sensitivity and specificity at different thresholds for SNP selection. We track power of different methods along different combinations of sensitivity and specificity parameters that lead to specific values of the area under the curve (AUC), which is an overall summary of the ability of the underlying score to discriminate between causal and non-causal variants (see Figure 4).

### Properties of the first- and second-order approximations

We conduct extensive simulation studies to evaluate accuracy of the proposed power calculations for variance component tests in comparison to exact theoretical methods that require specification of effect-sizes of individual variants. Here we focus on the SKAT test statistics as a representative of variance component tests and in Supplemental Materials we present results for the burden test statistic as a representative of sum-based tests. For each fixed combination of the size of a region (*J*) and the total variance explained (*TEV*), the two key parameters that determine the approximate power of the as SKAT test, we simulate various possible values of allele frequencies and effect sizes for individual markers *EV*_*j*_. Then the power based on the first- and the second-order approximations averaged over empirically derived distribution of allele frequencies is compared with power based on exact theoretical calculations averaged over distribution of allele frequencies and distribution of effect sizes for individual markers *EV*_*j*_ (see Figures 1 and 2, Appendix D for more detail).

**Figure 1.**
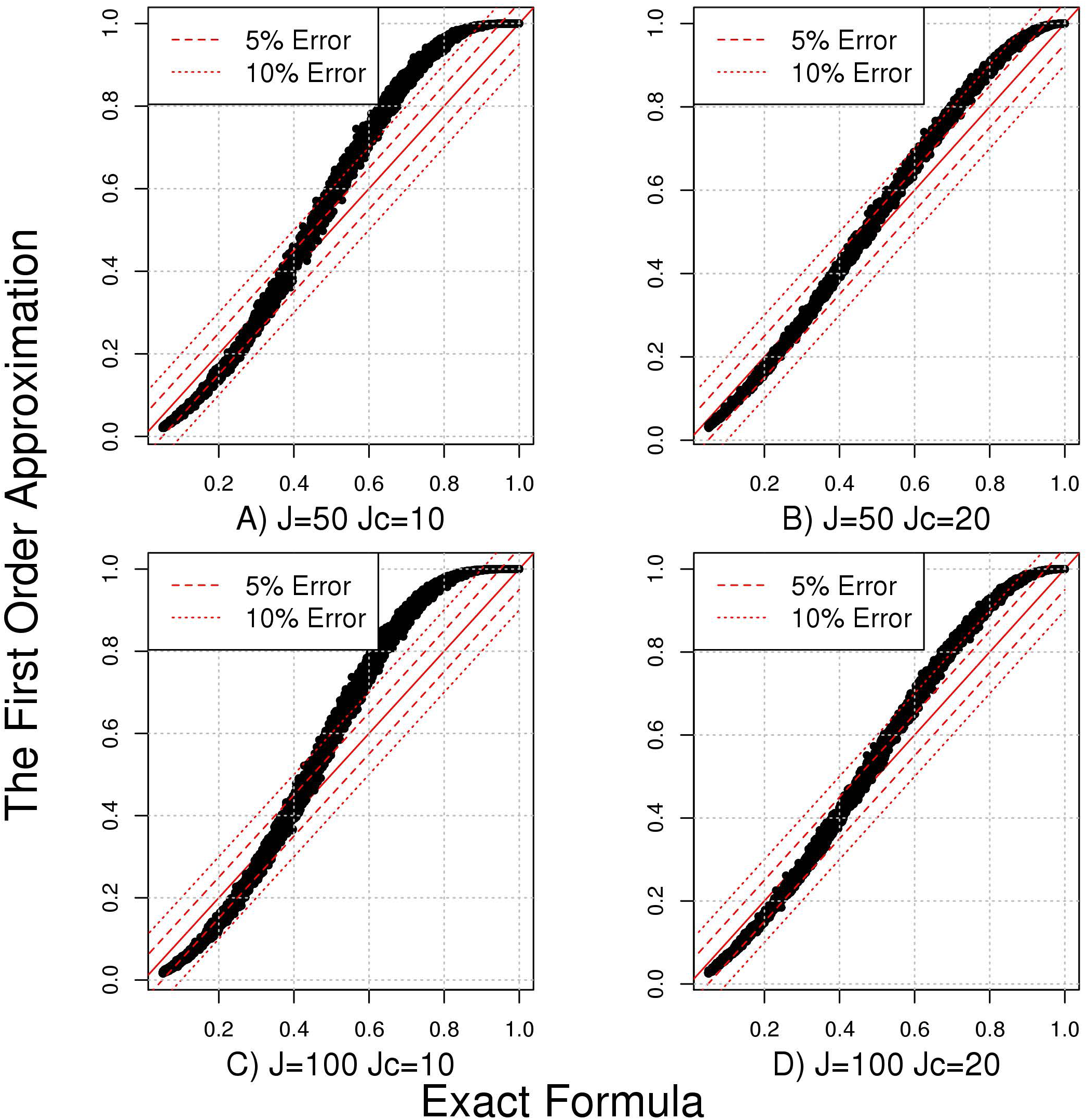
Evaluation of the accuracy of the first order approximations under simulation scenario S1 (MAF-independent EV). Exact Formula represents estimated average power using exact theoretical formulas for the SKAT test statistic. The First Order Approximation represents estimated average power using the first order approximation for the SKAT test statistic.

**Figure 2.**
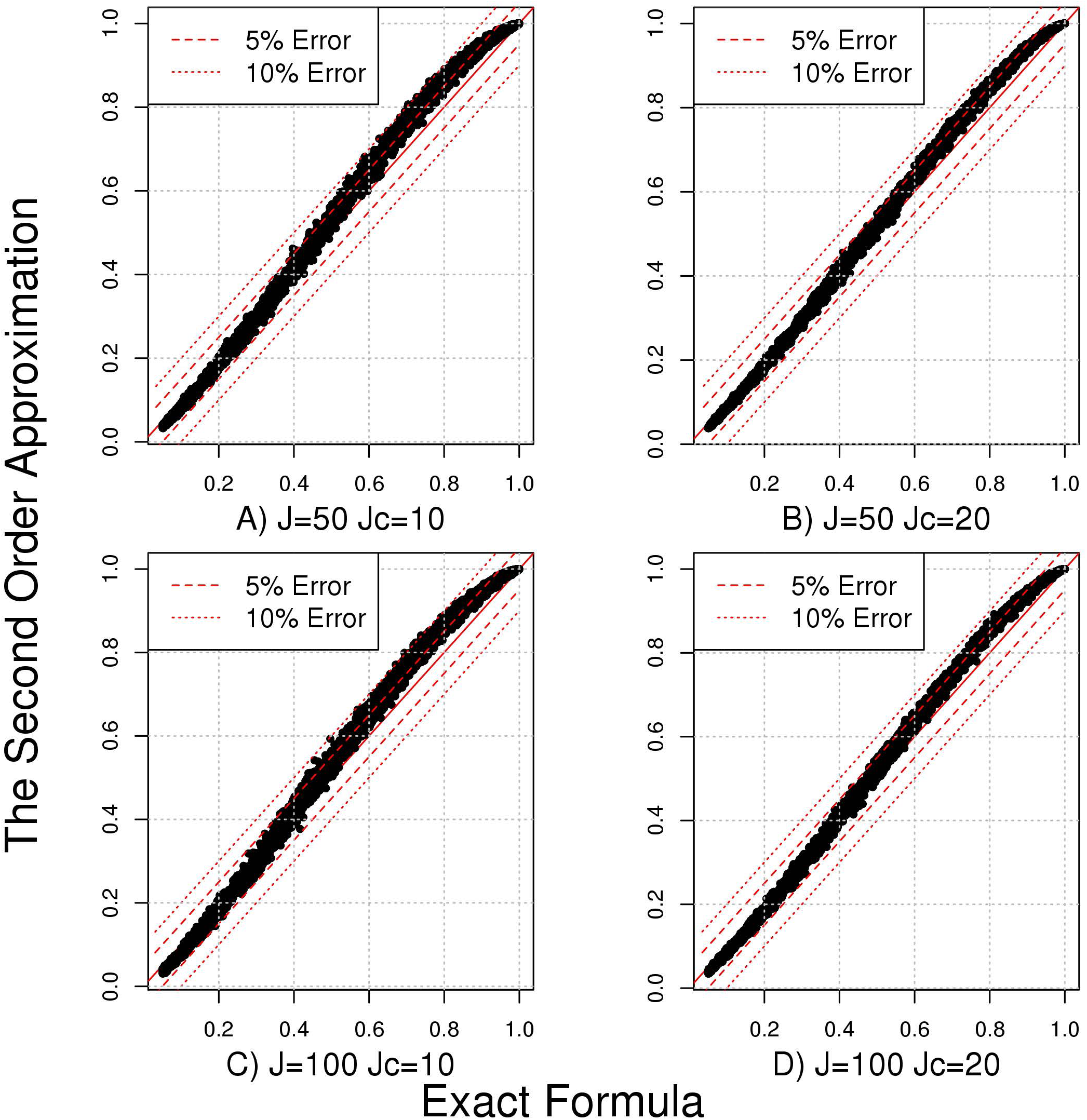
Evaluation of the accuracy of the second order approximations under simulation scenario S1 (MAF-independent EV). Exact Formula represents estimated average power using exact theoretical formulas for the SKAT test statistic. The Second Order Approximation represents estimated average power using the second order approximation for the SKAT test statistic.

We consider three types of simulation scenarios: S1 (″MAF-independent EV″) assumes that coefficients of explained variations (*EV*)is independent of MAF; S2 ('MAF-independent *β*_*j*_′) assumes that size of genetic effect (*β*), measured in the unit of per copy of an allele (*β*^2^ = *EV*/2*MAF* (*1-MAF*)) is independent of MAF and S3 ('MAF-log-dependent *β*_*j*_′) assumes that genetic effect is related to MAF through *log* 10 function (as defined in Table S1). For each type of simulation scenario, we estimate the power for a locus of size *J*=50,100,200 and 400 with the number of underlying causal variants *J*_*c*_ *=* 10,20,30 and 50. In Appendix D, we describe simulation mechanisms in detail and we summarize simulation models and parameters required for each method in Table S2.

### Bounds on variation explained by a causal locus

In Table S3, we provide key parameters that summarize a variety of recently published association studies of rare variants^15; 18; 21-23; 25-29^. Typically, studies on Human Exome BeadChip (Exome Chip) had larger sample sizes then studies on sequencing platform; however, the latter covered a much smaller number of rare variants. We use our mathematical framework to obtain bounds on genetic architecture implied by the results from two of the largest studies, one of which studied educational attainment with exome sequencing and the other studied blood pressure employing exome chip ^26; 29^ (see 5^th^ and 9^th^ rows of the Table S3).

To estimate genetic bounds from the results of study on educational attainment, we estimate the probability of no discoveries (*m*=0) given a genetic model from formula (2) by using the sample size *N* = 14,000, the number of sequenced individuals. We assume that analysis was conducted by the SKAT statistic with a gene as a unit. We use publicly available EXaC database^47^ to obtain empirical distributions for number of rare variants in a gene (*J*_*k*_) and vector of MAFs ***p***_*k*_ across exome (see Figures S9). To ensure validity of asymptotic power formulas, we assume that MAF of rare variant ranges between 0.0001 and 0.01 (e.g. no singletons and doubletons). Lastly, we set Type 1 error threshold 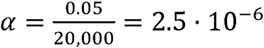.

To estimate genetic bounds from the study of blood pressure outcome, we calculate probability of at most three discoveries (m=3) under a sample size of *N =*140,000 to match number of discoveries reported and number of individuals genotyped in this study (see 9^th^ row of the Table S3). Similar to the first study, we assume that analysis was conducted by the SKAT statistic with a gene as a unit. Empirical distributions for number of rare variants in a gene (*J*_*k*_) and vector of MAFs ***p***_*k*_ are obtained also from EXaC database (see Figures S10). In contrast to the previous study, here we did not put any lower bound for MAFs. Lastly, we use the same Type 1 error threshold *α* = 2.5.10^-6^. The average numbers of rare variants per gene (*J*) were 35.5 and 13 in the studies of educational attainment and blood pressure, which used exome sequencing and Exome Chip platforms, respectively. We provide empirical distributions and other key parameters in Figures S9 and S10.

For each combination of the number underlying of causal loci (K) and parameters of effect size distributions, we calculate probability of less than or equal to m discoveries using formula (2) with study specific parameters. Expectations in (2) are estimated using 100,000 Monte Carlo simulations. In this report, we calculate power of the SKAT test statistic under the assumption of independence between proportion of variations explained and MAF of individual variants. Results for other genetic architecture are also discussed and presented in the Supplemental Materials.

### Effects of variant filtering on power of aggregated tests

We consider two values of a size of a locus *J*=50,100 and two values of a number of causal variants in a locus, *J*_*c*_ = 10,20. Initial values of *TEV* are selected so that power of the SKAT test is equal to 40% at Type 1 error α=0.05/20,000-2.5·10^-6^ and sample size *N*=10,000. For every combination of sensitivity and specificity, we estimate average power for the SKAT and burden tests. For this study, we also assume independence between proportion of variations explained and MAF for the individual variants. Results for other genetic architectures are presented in the Supplemental Materials.

## Results

### Properties of the First- and Second-Order Approximations

We evaluate accuracy of first- and second-order approximations compared to exact power calculations of the variance component test under variety of genetic models (see Table S2). The first order approximations match exact calculations better as the number of causal variants in a locus *J*_*c*_ increases (Figure 1). Particularly, we observe that with more than 20 causal variants in a locus (*J*_*c*_ ≥ 20), difference in power between those two methods is small regardless of the total number of variants in a locus *J*=50,100 (see Figure 1 B and D and Figure S1). Similar conclusion holds also for very large loci *J*=200,400 and other relationships between genetic effect and MAF (see Figures S2, S3 and S4). With lower number of causal variants in a locus (e.g. *J*_*c*_ =10), we observe upward bias in first-order estimates when exact power is high and downward bias when exact power is low (see Figure 1 A and C).

We observe that the second-order approximation is more accurate at estimating the exact power (Figure 2). Now, difference in power between approximate and exact calculations is small even when the number of causal variants in a locus is small, (see also Figures S5-S7). Overall, our simulations demonstrate that the first-order approximation accurately estimates exact power when number of causal variants in a locus is not too small. However, if the number of causal variants in a locus is small, then the first order approximation may produce biased results. On the other hand, the second-order approximation estimates exact power more accurately regardless of underlying generic architecture, but it requires specification of an additional parameter, namely the number of causal variants in a locus (*J*_*c*_).

As for linear statistic, we observe that the accuracy of the proposed approximations does not drastically depend on number of causal variants in a locus (Figure S8). However, the accuracy depends on the variation in SNP specific coefficients of variations (see Appendix A).

### Bounds on Effect Size Distribution

Genome-wide power analysis of the educational attainment study^26^ (Figure 3 A), which implemented whole exome sequencing, shows implausibility of models that correspond to a small number of underlying causal loci explaining significant total phenotypic variance. For example, the probability of observing no discovery is less than 5% under genetic models that involve less than 250 loci to explain a total of 20% or more phenotypic variation. If we assume independence between genetic effects and MAFs across variants, a scenario under which SKAT test has higher power, then even larger number of loci will be needed to explain the same total variance (see Figure S11 A). Genome-wide power analysis of the study of blood pressure^29^ (Figure 3 B), which is much larger in sample size but implements the Exome Chip platform, provides very sharp bound on the relationship between number of underlying causal loci and total heritability explained by the underlying variants. We estimate, for example, at least 6000 causal loci will need to be involved if the variants included in the study could explain 20% of phenotypic variance of blood pressure. Identical to results for the WES study, genetic bound is even sharper if independence between MAFs and genetic effects is assumed (see Figure S11 B).

**Figure 3.**
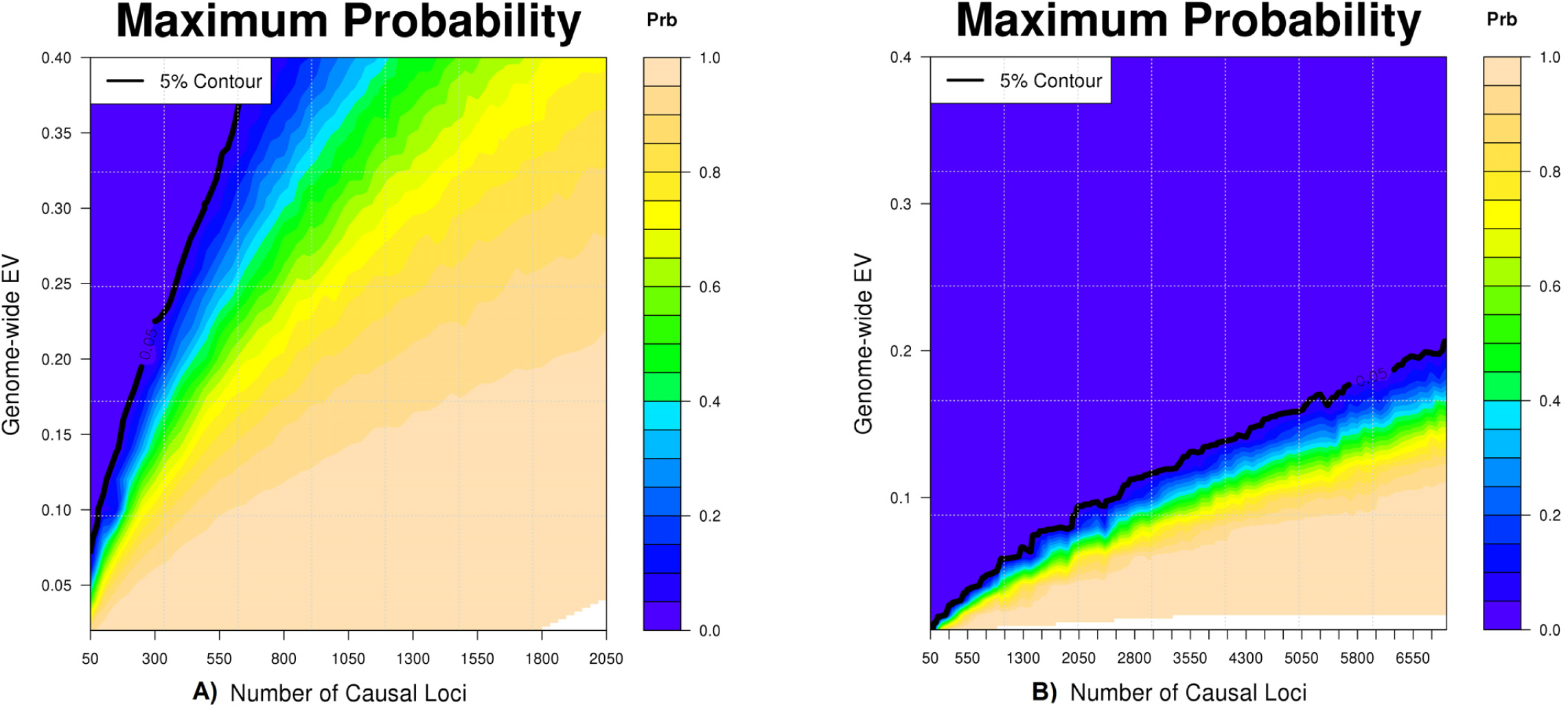
Bounds for genetic architecture based on results reported in studies of education attainment (EA) and blood pressure(BP). Panel (A) shows maximum probability of observing no discoveries in the EA study, which used whole exome sequencing platform, as a function of the number of underlying causal loci K and the total variation explained by them with a sample size of 14,000. Panel (B) shows the maximum probability of observing three statistical significant discoveries in the BP study, which used exome chip, as a function of the number of underlying causal loci K and the total variation explained by them with a sample size of 140,000. In both cases, it’s assumed gene-based tests have been performed using the SKAT test statistics at the level of α =2.5·10^-6^. Probabilities are estimated by (2) and assumption of independence between MAF and EV. The black line shows approximate contours (bounds) corresponding to probability of 5%.

### Effects of Apriori SNP Screening on Power of Aggregated Test

Apriori SNP selection does not improve the power of variance-component test substantially (e.g. by 10%) unless the underlying algorithm has very high accuracy to discriminate between causal and non-causal SNPs (AUC between 80-90%) (see Figure 4). In contrast, power for sum-based test can improve substantially with more modest discriminatory accuracy of the SNP selection algorithm (AUC between 70-80%). Further, we observe the roles of sensitivity and specificity are not symmetric on power of these tests. For both tests, substantial improvement of power is possible only if sensitivity is at the minimal 30-40%. On the other hand, substantial improvement in power is possible with fairly poor specificity (e.g. about 20%) as along as sensitivity is high (e.g. 90%). We observe similar results in studies with different genetic architecture and large number of SNPs in a locus (see Figures S12-S13).

**Figure 4.**
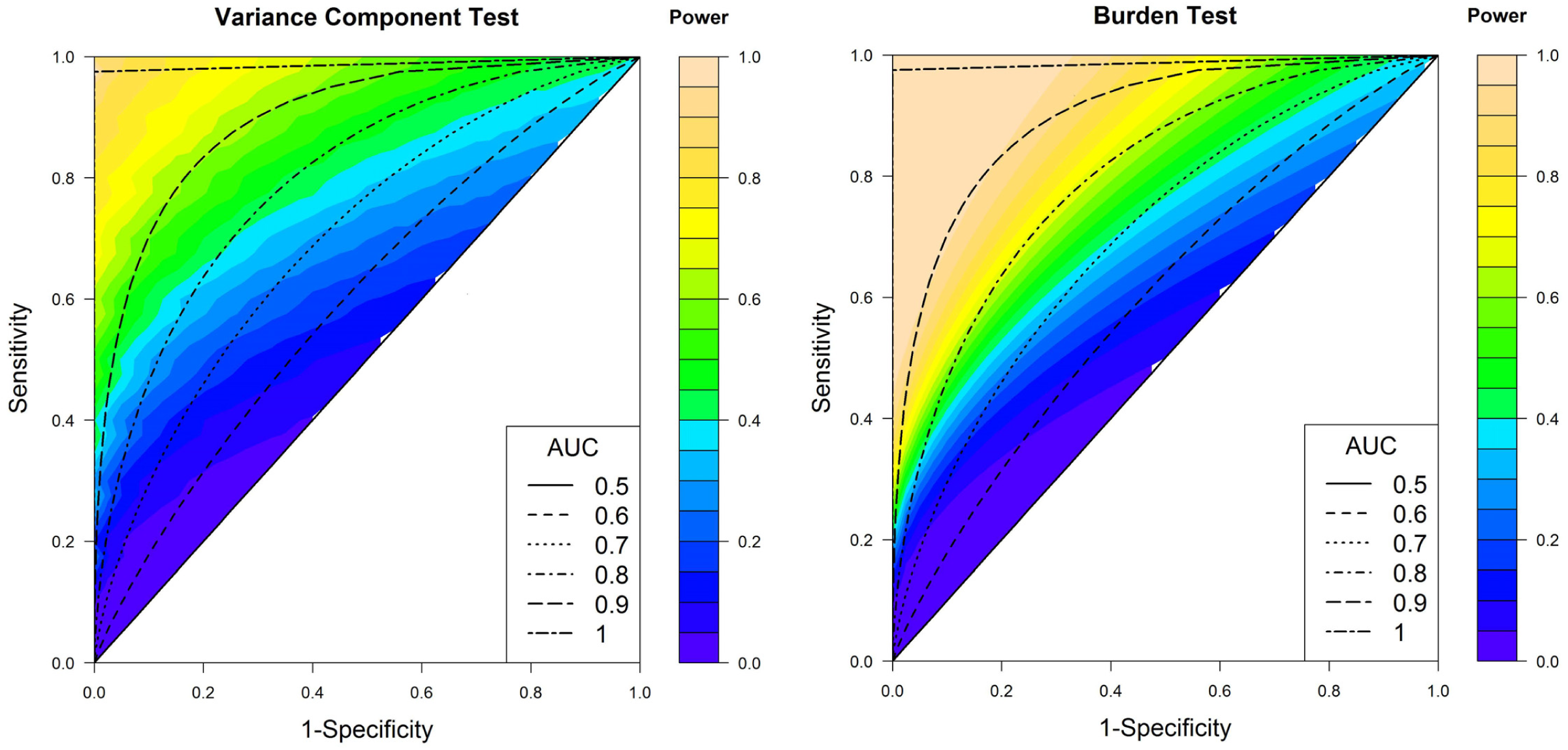
Effects of sensitivity and specificity for apriori variant screening on the power of variance component and burden tests under simulation scenario S1 (MAF-independent EV). Number of variants in a locus is set to **J=50** and number of causal variants to **J_C_=10**. The setting corresponds to a baseline power (i.e. if all variants were included in the study) of 40%, and 36% for variance component and sum-based tests, respectively.

## Discussion

Although large GWAS of low frequency and rare variants are now becoming increasingly feasible due to technological advances, the likely yield of such studies in the future remains uncertain as studies conducted to date have only reported limited number of findings^15-30^. For studies of common variants, which have mostly relied on association testing at the level of individual variants, we and others have shown that the yield of genome-wide association studies critically depend on distribution of phenotypic variances explained by individual variants across the genome^4; 7; 52^. For studies of rare variants, it has been suggested that tests for genetic associations be performed at an aggregated level by combining signals across multiple variants for powerful detection of underlying susceptibility loci^30; 33-35; 44^. In this report, we show that how power for some of these more complex tests critically relates to total genetic variances explained by multiple variants within a locus. Based on such power calculations, we assess bounds on distributions of locus-level genetic variances that are consistent with limited findings reported in current studies. Further, based on these simplified power calculations, we evaluate potential for improving power for aggregated tests by pre-selection of likely causal variants based on functional/annotation information.

Power analysis of current studies of large sample sizes may provide important bounds on genetic architecture of the underlying traits. Our analysis suggests that rare variants investigated in current studies could explain significant fraction of heritability of the underlying traits only under highly polygenic models in which causal variants are distributed over hundreds or even thousands of different genetic loci. These results are intuitive given that if a relatively small number, e.g. a few dozens, of genetic loci could explain a substantial fraction of heritability of these traits, then at least some of these loci will be detected by the sample size achieved so far in the current studies.

A number of rare variant studies that have conducted both individual-variant and aggregated tests have detected more genetic loci using the former than the later approach^18; 23; 28; 29^ (see Table S3). The analytic formula we propose for calculating probability of certain number of discoveries under various models for genetic architecture can also be applied for single-variant tests. Genetic bounds based on the results from single SNP analysis for the same two studies also show that only under highly polygenic architecture the variants included in these studies can explain substantial fraction of heritability of the underlying traits (see Figures S14). These genetic bounds based on single SNP analysis are also consistent with corresponding genetic bounds from gene based analysis and assumption of clustering of multiple causal variants per causal locus. Large studies with more accurate estimates of genetic bounds will provide additional information on degrees of clustering of multiple rare variants within causal locus.

A variety of studies have studied genetic architecture of common variants by characterization of underlying heritability, number of susceptibility variants and effect-size distributions^4-11; 32; 53-57^. All of these studies consistently point toward a highly polygenic model where disease etiology may involve thousands or even tens of thousands common susceptibility variants, each conferring only a modest association, but in combinations they can explain substantial phenotypic variations. Some recent studies have reported that low frequency and rare variant studies have the potential to explain significant fraction of heritability for selected traits^18; 58; 59^. Further insights into genetic architecture of these traits can be obtained by comparing observed number discoveries in these studies with those from simulated studies under different models for genetic architecture^31; 60^. The proposed analytic framework provides an alternative fast and simple way of evaluating expected discoveries for a large variety of genetic models and quantification of their plausibility given results from a given study.

Power calculations for aggregated tests with selected subset of variants point towards challenges for use of functional and annotation information for pre-screening. Overall, it appears that pre-selection of variants can significantly improve the power of aggregated tests only if the underlying functional/annotation information have fairly high accuracy to discriminate (AUC > 70-80%) between causal and non-causal variants for the underlying disease of interest. In particular, the algorithm should be highly sensitive to capture the underlying causal variants for a disease. Use of too stringent criterion for variant selection may increase specificity but will lead to decreased sensitivity and hence could lead to loss of power in aggregated tests. More empirical studies are needed to assess the impact of variant selection on power of aggregated tests.

Sophisticated imputation algorithms^12; 13^ and increasing sample size of reference datasets^12^ are allowing imputation of low frequency and rare variants with increasing accuracy. Many association studies are now being conducted based on imputation in existing large GWAS. A limitation of our method is that it currently cannot account for imputation accuracy, which is expected to reduce with decreasing allele frequency. At the level of individual variants, it is possible to characterize reduction of power based on formula for effect-size attenuation due to imputation^61^. Further studies are needed to understand impact of imputation on aggregated tests encompassing variants of different allele frequency spectra. In this report, we have illustrated application of the framework in exome-based analysis where aggregated tests can be applied across largely non-overlapping genes. For whole genome sequencing studies, where aggregated tests may be applied in a sliding window fashion^18; 22^, more work is needed for genome-wide power calculations in terms of underlying models for genetic architecture.

In conclusion, in this report we provide simple analytic approaches to power calculations for rare variants association tests at the levels of individual loci and whole genome in terms of a few key parameters of the underlying models for genetic architecture. These methods, which we implement in a Shiny application in R, will provide useful design tools for planning next generation genome-wide association studies.

## Appendix A Approximation to Sum-based Tests

The non-centrality parameter of a linear statistic is 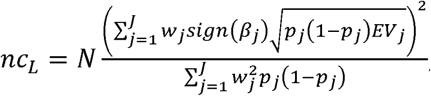, where *J* is a number of SNPs in a locus, *p*_*j*_ is MAF, *EV*_*J*_ is a proportion of phenotypic variation explained by SNP *j* and *w*_*j*_ is MAF-based weight. Let *J*_*D*_ is number of variants with positive effect and *J*_*p*_ is number of variants with negative effect. Non-centrality parameter *nc*_*L*_ is rewritten as

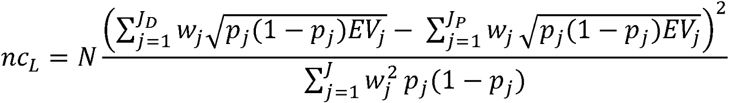

With sufficiently large *J*_*D*_ and *J*_*p*_, sums in numeration of non-centrality parameter can be approximated as expected values and hence *nc*_*L*_ is approximated by

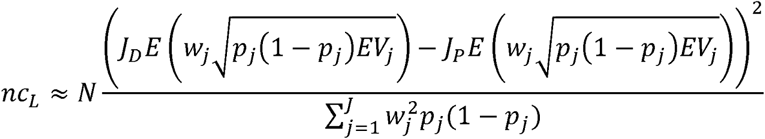

Under low variation in *p*_*j*_ and *EV*_*j*_, previous equation is further simplified

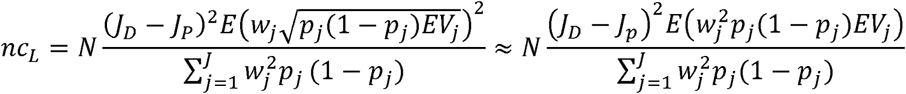

Under independence between *p*_*j*_ and, 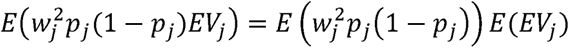 and *nc*_*L:*_ is finally simplified to

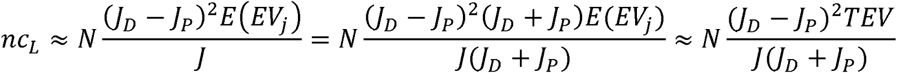

where *EV* is proportion of variation explained by SNPs.

## Appendix B First- and Second - Order Approximation to Variance Component Tests

We assume that genetic effect *β*_*j*_ *= f(p*_*j*_*) e*_*j*_ depends on MAF *p*_*j*_ through some continuous function f() and random effects *e*_*j*_ Three common relationships between genetic effect and MAF are 1) coefficient of explained variation by a SNP *EV*_*j*_ is independent of (e.g.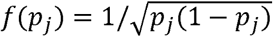, 2) genetic effect *β*_*j*_ is independent of *p*_*j*_ (e.g. *f*(*p*_*j*_) = 1) and 3) genetic effect *β*_*j*_ is related to *p*_*j*_ by *f*(*p*_*j*_) = *log*_10_(*p*_*j*_). Coefficient of variation explained by a SNP is *EV*_*j*_ = 2*p*_*j*_(1 − *p*_*j*_)*f*(*p*_*j*_)^2^*e*_*j*_^2^. For considered small values of *p*_*j*_, *EV*_*j*_ can be approximated as linear function of 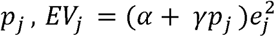. Under relationship 1, α and γ are equal to and 1 and 0. Under relationship 2, α and γ are equal to 0 and 1. Under relationship 3, *α* and *γ* are solutions to 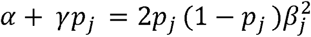 with *p*_*j*_ equals to lower and upper MAF bounds.

Power of variance-component statistic can be approximated by a chi-square distribution with the first four cumulants specified as

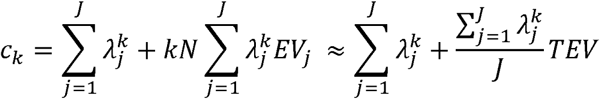

The first order approximation estimates sums 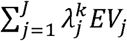 in cumulants as functions of 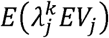. With three proposed relationships between MAFs and *EV*_*j*_, we obtain following approximations

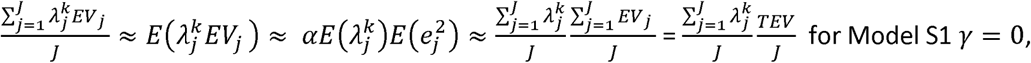

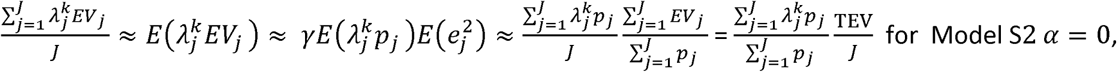

and

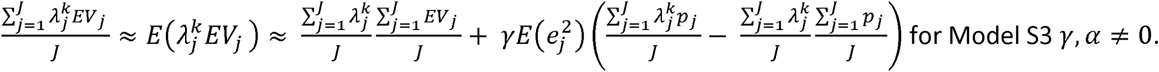

Parameter 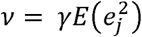 represents average change of in due to one unit change in MAF *p*_*j*_. By plugging 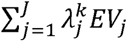 into cumulants *c*_*k*_, we obtain the first order approximations presented in Table S1.

The second order approximation is almost identical except instead of *J* in we use *J*_*c*_, number of causal variants in a gene (see Table S1)

## Appendix C Probability of M Discoveries in a Genome-wide Study with an Aggregated Level Test

Let *pow*_*k*_ *= pow*(*α,N,TEV*_*k,*_*J*_*k,*_***p***_*k*_) is power of an aggregated-level test for the causal locus *k*. Probability of M discoveries in a study with *K* underlying loci is

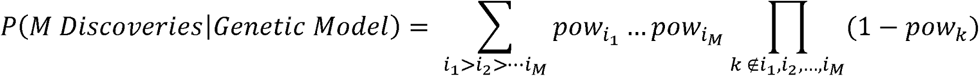

Under assumptions *M*≪ *K* (is large),

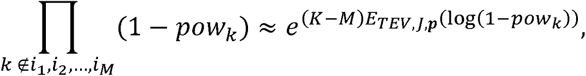

for any set of indices (*i*_1_,*i*_*2*_,…,*i*_M_). Similarly, the sum 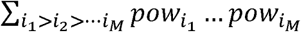 can also be approximated as

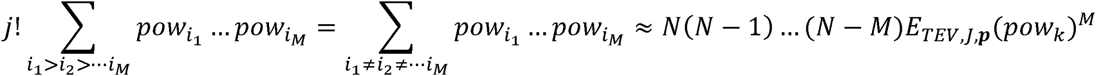

while assuming independence between loci. As result, the probability of discoveries can be approximated by

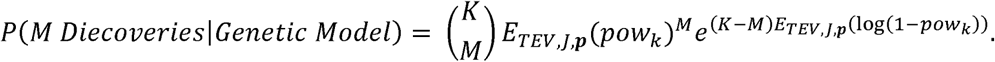

## Appendix D Simulation Procedure

For each simulation set-up, we fixed a value of the total number of variants *J* and the number of causal variants in a locus *J*_*c*_, we estimate average power of two methods over a wide range of *TEVs* (see Table S2). For each value of *TEV* and *J*, we repeatedly generate *J* MAFs from Gamma distribution with shape and scale parameters from ExAC database^47^ and calculate average power over 1000 such replications under the first-order approximations. The average power under exact theoretical calculations is also obtained based on 1,000 replications. However, here, addition to a vector of MAFs, we generate coefficients of explained variations *EV*_*j*_ for all causal variants *J*_*c*_. For three genetic models S1-S3 (see Table S1 and S2), we use different strategies for generating SNP specific coefficients of explained variations. For the scenario S1, total coefficient of variation *TEV* is randomly split into *J*_*C*_ SNP specific coefficient of variations *EV*_*j*_. For the scenarios S2, we calculate *β*^2^,a MAF adjusted average effect of a variant and for the scenario S3, we calculate normalizing constant *C* as presented in Table S2. Then SNP specific coefficient of variations *EV*_*j*_ are generated using MAFs and *β*_*j*_^2^. For example, we generate 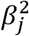 from 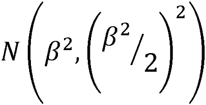 for the scenario S2 and recalculate 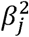 from *Clog*_10_(*p*_*j*_)^2^for the scenario S3.

Lastly, we scale back so that total variation explained by causal variants is equal to.

## Supplemental Data

Supplemental Data include three tables and fourteen figures and can be found with this article online at

## Acknowledgments

This study utilized the computational resources of the NIH HPC Biowulf cluster. AD is supported by the National Cancer Institute Intramural Research Program.

### Web Resources

The URLs for data presented herein are as follows:

*PAGEANT*: https://andrewhaoyu.shinyapps.io/shinyapp/

